# Same-section spatial metabolo-transcriptomics using Stereo-meta-seq reveals DHA-driven kidney maturation

**DOI:** 10.64898/2026.06.16.732541

**Authors:** Roan H. van Scheppingen, Baihua Wu, Yumeng Zhang, Sytske Schep, Ulli Winter, Fengxiang Wang, Yilong Zou, Susana M Chuva de Sousa Lopes, David Silver, Iulia Sidorov, Martin Giera, Bernard M. van den Berg, Cathelijne W. van den Berg, Lijian Hui, Ton J. Rabelink, Gangqi Wang

## Abstract

A central unresolved question in developmental biology is whether local metabolites merely accompany, or actively instruct, tissue maturation. Addressing this question requires direct spatial coupling of metabolic states with genome-wide transcriptional programs *in situ* at high spatial resolution, which existing approaches do not readily achieve. Here, we introduce Stereo-meta-seq, a workflow that integrates quantitative MALDI-MSI with Stereo-seq spatial transcriptomics within a single tissue section. A conductive adapter was designed to overcome the electrical incompatibility of non-conductive Stereo-seq chips with vacuum MALDI platforms, improving efficiency of MSI detection that preserves RNA integrity. MALDI laser-ablation marks are retained in downstream Stereo-seq data and serve as intrinsic fiducials for direct co-registration at 10 μm or 20 μm resolution, enabling fine grained spatial metabolite-transcript integration. Applying Stereo-meta-seq to human kidney development, we uncover selective enrichment of docosahexaenoic acid (DHA) in maturing proximal tubules. Functional studies in human kidney organoids demonstrate that DHA activates PPARα-and HNF4α-driven transcriptional programs and promotes proximal tubule maturation *in vitro* and after transplantation *in vivo*. These findings identify lipid metabolism as an instructive regulator of human nephrogenesis and establish Stereo-meta-seq as a practical platform for dissecting metabolite–gene coupling and tissue heterogeneity *in situ*.

## Introduction

Tissue development is governed not only by gene-regulatory networks but also by spatially organized metabolic programs that actively shape cell fate transitions. Metabolites can regulate signaling pathways, transcription factor activity and chromatin state, thereby coupling nutrient availability to developmental gene expression programs^1, 2^. In pluripotent stem cells, intracellular α-ketoglutarate has been shown to sustain self-renewal by promoting a chromatin state compatible with naïve pluripotency, whereas glycolysis-dependent acetyl-CoA production supports histone acetylation during the early transition out of pluripotency^3^. More recently, spatially resolved analyses of mammalian gastrulation revealed cell-type- and stage-specific waves of glucose metabolism that guide lineage specification and morphogenesis *in vivo*^4^. Together, these findings indicate that metabolism is not merely a downstream consequence of developmental state, but an instructive layer of regulation that must be measured in its native spatial context, motivating technologies capable of jointly resolving metabolites and transcriptomes within intact tissues^5^.

Spatially resolved molecular profiling has transformed tissue biology by enabling the study of cellular states within their native anatomical context. Among these technologies, spatial transcriptomics has become a powerful framework for mapping genome-wide gene expression across intact tissues, revealing spatial organization, cellular interactions, and regional heterogeneity that are inaccessible to dissociative single-cell approaches^6, 7^. However, transcriptional states do not fully capture the biochemical outputs that directly shape phenotype, particularly in processes driven by nutrient environment and metabolic remodeling. MALDI-MSI provides label-free spatial detection of metabolites, lipids, peptides and other molecular classes directly from tissue sections and has emerged as a core platform for spatial metabolomics^8^. Integrating MALDI-MSI with spatial transcriptomics therefore offers a direct route to connect molecular regulation with biochemical phenotype *in situ*. Vicari *et al*. (2024) combined MALDI-MSI with Visium on the same tissue section, demonstrating that same-section acquisition can retain analytical performance in both modalities^9^. Same-section integration of MALDI-MSI with Xenium showed that MALDI laser-ablation marks can be used as fiducials for direct alignment, although prior MSI reduced downstream transcript counts per cell by approximately 30%^10^. In parallel, alternative same-section strategies such as MALDI-ISH MSI have further underscored the growing interest in joint metabolite-RNA mapping^11^. Nevertheless, current approaches remain constrained by one or more of the following limitations: low spatial resolution, targeted transcript panels, modality-dependent registration complexity or limited quantitative capability in MSI.

Stereo-seq is particularly attractive for addressing these limitations because it offers sequencing-based, transcriptome-wide spatial readout with high physical resolution and large field of view^6^. However, direct integration of Stereo-seq with vacuum MALDI imaging platforms presents a practical obstacle: Stereo-seq chips are not intrinsically optimized as conductive MALDI devices. In routine MALDI-MSI workflows, electrically conductive supports such as indium tin oxide (ITO)-coated slides are generally recommended, whereas poorly conductive surfaces can compromise ion extraction and image quality because of charging effects. Accordingly, custom holders, conductive coatings, conductive tapes, and conductive adhesive films have all been explored to restore signal quality in otherwise poorly conductive samples^12, 13^. However, these solutions were not developed for seamless coupling to genome-wide, sequencing-based Stereo-seq, and same-section MALDI workflows can compromise RNA integrity, resulting in a measurable loss in downstream transcriptomic quality.

Here, we present Stereo-meta-seq, a same-section spatial multi-omics workflow that integrates MALDI-MSI-based metabolite mapping with Stereo-seq spatial transcriptomics. By addressing the compatibility challenge between non-conductive Stereo-seq chips and MALDI imaging, this approach enables direct spatial integration of metabolites and genome-wide transcripts at high spatial resolution on the same tissue section. Applied to human kidney development, Stereo-meta-seq identifies selective enrichment of docosahexaenoic acid (DHA, C22:6n3) in maturing proximal tubules and links this metabolic feature to transcriptional programs associated with epithelial maturation. More broadly, this workflow provides a practical framework for dissecting metabolite–gene coupling and tissue heterogeneity *in situ*.

## Results

### Stereo-meta-seq workflow

A key innovation in our approach is the design of a conductive adapter that enables robust MALDI imaging of non-conductive Stereo-seq chips on a Bruker rapifleX mass spectrometer and improves acquisition efficiency on timsTOF fleX MALDI-2. This design stabilizes electrical coupling at the sample interface, supports high-quality ion extraction on transcriptomics-compatible substrates, and preserves MALDI laser-ablation marks as intrinsic spatial landmarks for direct co-registration with downstream Stereo-seq data (Figure 1A). As a result, our workflow enables direct metabolite and genome-wide transcriptome integration at 10 μm or 20 μm resolution on the same tissue section.

**Figure 1.**
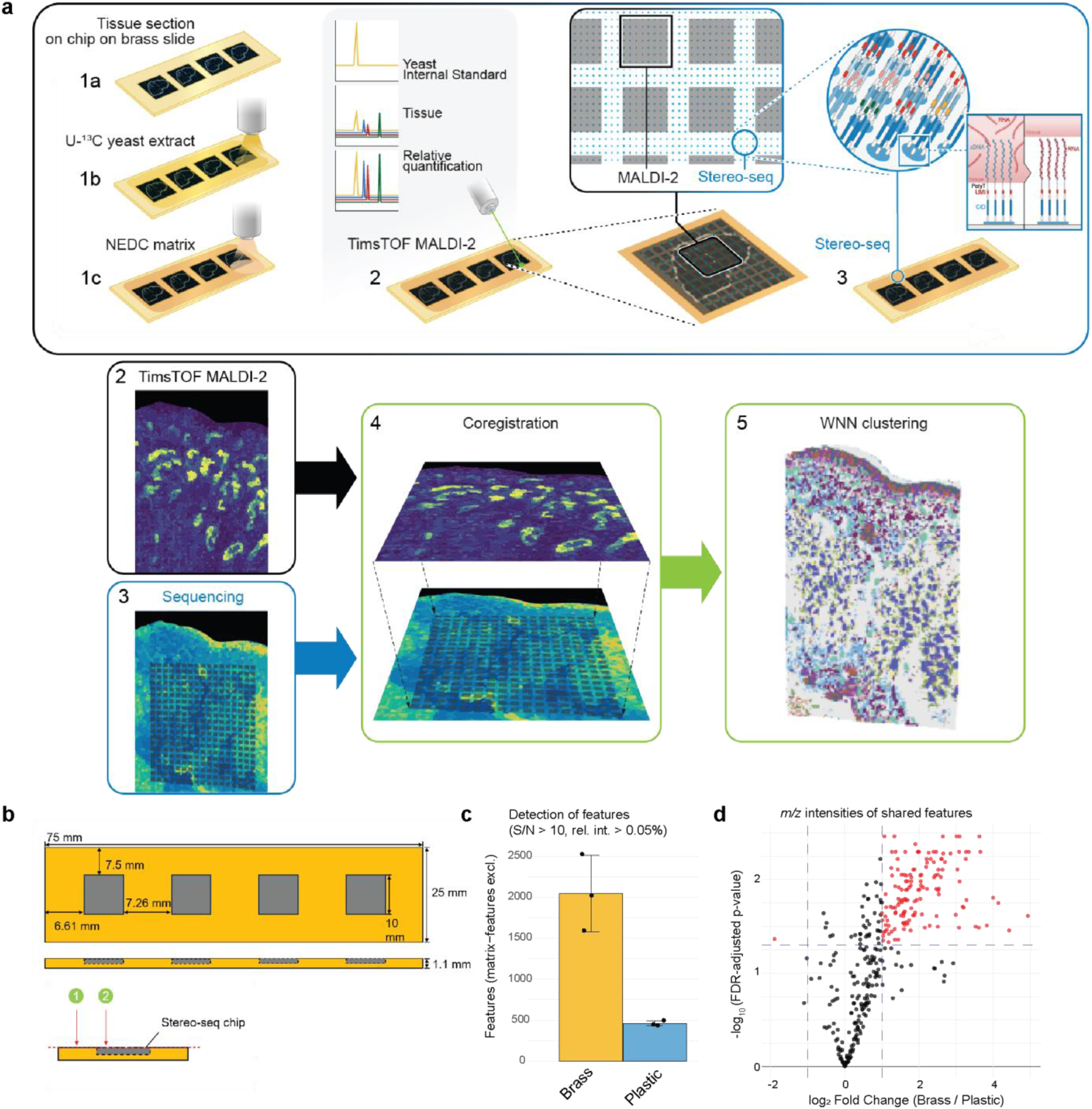
Stereo-meta-seq achieves single slide multi-omics measurements through higher sensitivity during MSI acquisition. (A) Schematic overview of experimental workflow of Stereo-meta-seq. (B) Dimensions of the custom-made brass docking slide. (C) Quantification of the number of features reaching signal-to-noise > 10 and a relative intensity > 0.05% in qMSI per material. (D) Volcano-plot of fold changes of shared features between brass and plastic docking slides.

To enable same-section MALDI-MSI on Stereo-seq chips, the conductive brass adapter addresses two major determinants of MSI performance: electrical conductivity at the sample interface and precise alignment of tissue height with the surrounding slide surface (Figure 1B). Both factors are critical for stable ion extraction and high-quality spectral acquisition. The adapter embeds the non-conductive Stereo-seq chip within a conductive frame while keeping the tissue surface flush with the slide plane. An identically shaped plastic adapter was generated as a non-conductive control. On the Bruker rapifleX platform, the brass adapter produced MSI data comparable to those obtained from conventional ITO slides, whereas the plastic adapter failed to support usable signal acquisition (Supplementary Figure 1). We next evaluated both adapters on the Bruker timsTOF fleX MALDI-2 platform. Although this system is relatively tolerant of non-conductive substrates, the conductive adapter still increased the number of detected non-matrix spectral peaks by approximately four-fold under identical acquisition settings on the same kidney tissue (Figure 1C). Moreover, for spectral peaks detected with both adapters, the brass adapter consistently yielded higher peak intensities for most features (Figure 1D). These results demonstrate that the conductive adapter is essential for robust MSI acquisition on Stereo-seq chips and enables high-quality metabolic profiling while minimizing laser-induced tissue damage.

### Stereo-meta-seq generalizes across tissues

We first evaluated Stereo-meta-seq in mouse kidney and human fetal kidney tissues at 20 μm resolution with the timsTOF flex MALDI-2. For these experiments, we used 16 μm thick sections. Because MALDI-MSI detects metabolic features released from the laser-ablated tissue surface, excessive ablation can compromise downstream RNA capture. To preserve transcriptomic information while maintaining MSI performance, we optimized the acquisition method while using the conductive adapter and used section thickness sufficient to ensure that only part of the tissue volume was sampled during MSI. Under these conditions, Stereo-meta-seq enabled joint metabolomic and transcriptomic profiling within an effective 20 × 20 × 16 μm voxel. Combined with our previous developed spatial quantitative metabolomics method^14^, we can detect up to 198 metabolic features with pixel-wise normalization and up to 1004 gene features on average per voxel in the human fetal kidney.

To further assess the potential for near-single-cell analysis, we applied the workflow to mouse hippocampus using 10 μm resolution with the timsTOF flex MALDI-2 on 12 μm thick sections, enabling integrated metabolite and transcriptome readout within a 10 × 10 × 12 μm voxel (Supplementary Fig. 2). We then compared post-MSI Stereo-seq performance with previously reported spatial transcriptomics datasets^15^ and found that data quality after MSI remained within the range of published datasets generated without prior MSI, although some variation was observed across tissue types (Supplementary Figure 2D).

We next used human fetal kidney as a representative tissue to directly evaluate the impact of laser ablation on downstream spatial transcriptomic quality. Within the same section, regions subjected to MSI showed an approximately 20% reduction in transcriptomic counts relative to adjacent non-ablated regions (Supplementary Fig. 3A).

Despite this modest loss, post-MSI Stereo-seq data remained sufficient to resolve tissue heterogeneity and to reconstruct histology-like molecular patterns (Supplemental Figure 4A, 5A-D). Notably, during downstream Stereo-seq processing, ssDNA staining together with autofluorescence imaging independently revealed the ablated laser spots (Supplementary Fig. 3C). Moreover, laser-ablated positions could be identified in the Stereo-seq data itself through local reductions in captured mRNA signal, particularly in tissue-free regions and in areas of low cellular density (Supplementary Fig. 3D-F). These two sources of positional information provided intrinsic fiducials for co-registration of MSI and Stereo-seq data. Together, these results demonstrate that Stereo-meta-seq can be applied across tissue types to achieve high spatial resolution, same-section profiling of spatial metabolomics and spatial transcriptomics.

### Multimodal integration improves cell-state resolution during human nephrogenesis

We next focused on Stereo-meta-seq of human fetal kidney to investigate how cellular metabolism is linked to nephrogenesis. Weighted nearest neighbor (WNN)^16^ analysis combined metabolomic and transcriptomic features within shared spatial coordinates (Figure 2A). Multimodal clustering resolved 13 cellular populations spanning the nephrogenic trajectory and improved cell-state discrimination relative to either modality alone (Figure 2D; Supplementary Fig. 4). Nephron progenitor subpopulations were primarily defined by transcriptomic features, whereas proximal tubule identity was strongly reinforced by lipidomic features, highlighting the complementary contributions of the two modalities (Supplementary Fig. 4). Stereo-meta-seq also identified lipid-defined clusters that were not readily resolved by transcriptomics alone (Figure 2B–C).

**Figure 2.**
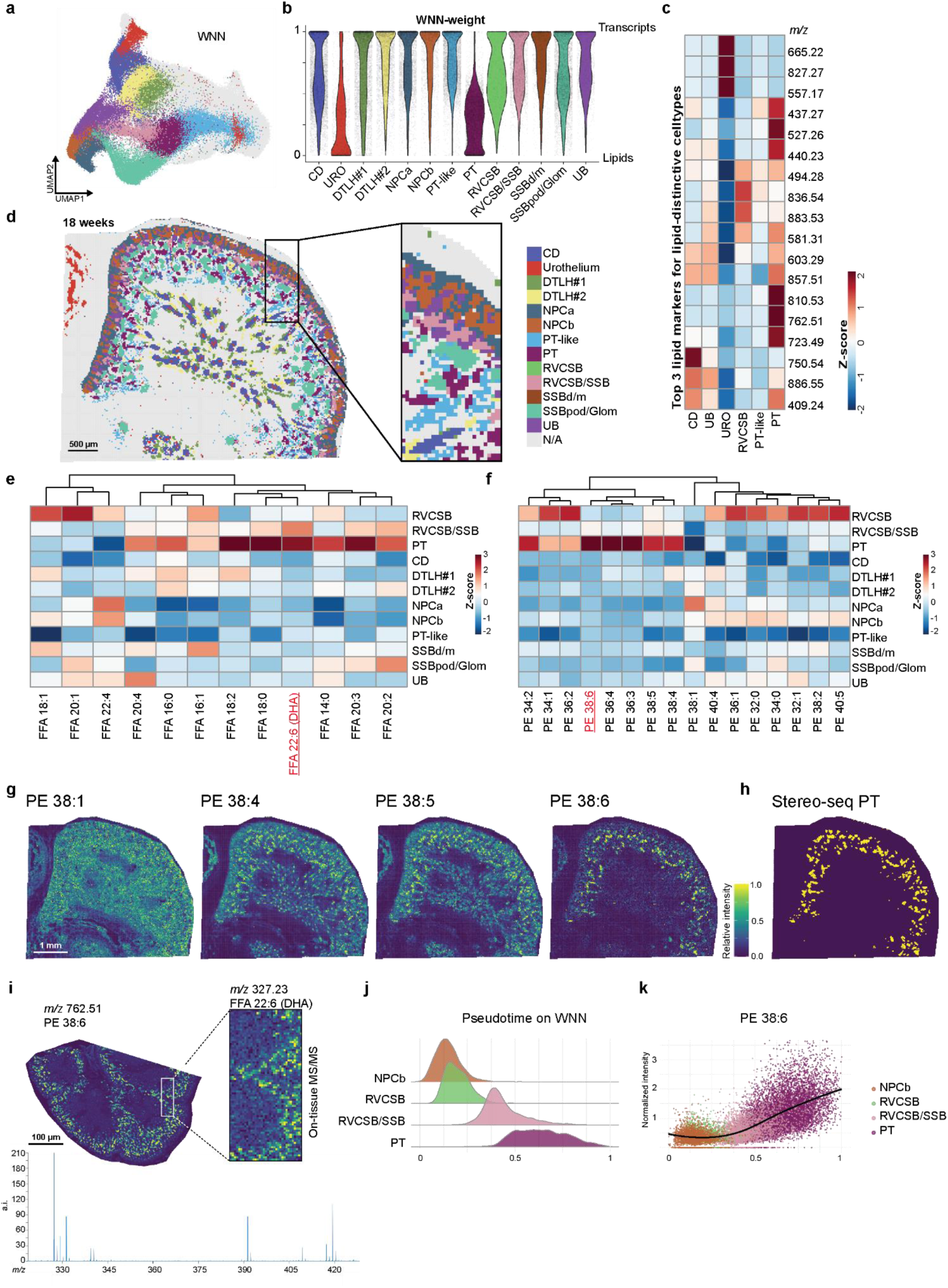
Stereo-meta-seq enables high-resolution cell type identification and reveals PE 38:6 as marker of proximal tubules. (A) Uniform Manifold Approximation and Projection (UMAP) after Weighted-Nearest Neighbor (WNN) clustering. (B) WNN-weight per cluster, a weight of 1 indicates that the transcriptomic modality was most deterministic. (C) Heatmap of Z-scores from the top three *m/z* features in clusters with low WNN-weight (lipidome-driven). (D) Representative image of a human fetal kidney with annotated clusters. (E) Heatmap of Z-scores of free fatty acids (FFA) detected using qMSI. (F) Heatmap of Z-scores of phosphatidylethanolamines (PE) detected using qMSI. (G) Spatial distribution of PE 38:*n* in the human fetal kidney. (H) Overview of cell population proximal tubules (PT) (yellow) in the human fetal kidney. (I) Tandem MS measurements of PE 38:6 (*left*) showing that PE 38:6 contains docosahexaenoic acid (DHA; *right zoom-in*). With a zoom-in of the MS/MS overview spectrum (*below*) (J) Pseudotime reconstruction from nephron progenitor cells (NPCb) towards PT. (K) Normalized intensity of PE 38:6 along pseudotime. **Abbreviations**; CD, collecting duct; URO, urothelium; DTLH, descending thin limb of the loop of Henle; NPC, nephron progenitor cell; PT, proximal tubule; RVCSB, renal vesicle/comma-shaped body; SSB, S-shaped body; pod, podocyte; Glom, glomerulus; UB, ureteric bud.

Notably, proximal tubules were difficult to delineate using Stereo-seq alone, in part because robust proximal tubule markers were not consistently resolved in pixel-based analyses of fetal kidney. This likely reflects the distinct tissue architecture of nephrogenesis: nephron progenitor and early nephron structures form dense cellular aggregates at the nephrogenic niche, whereas proximal tubules adopt a more dispersed tubular morphology. Differences in local cell density can therefore bias pixel-level transcriptomic assignment. For example, the proximal tubule maturation marker *CUBN* showed unexpectedly higher signal in S-shaped body regions (Supplementary Fig. 5E), and additional PT-like transcriptomic regions appeared without coherent tubular organization, suggesting misassignment or noise (Supplementary Fig. 4B). By contrast, multimodal integration substantially improved proximal tubule identification and separation, with lipid features, particularly *m/z* 762.51, that was pre-annotated as PE 38:6, providing strong discriminatory power (Figure 2C-F).

In addition to the detection of lipid features reflecting specific cell types, Stereo-meta-seq simultaneously profiles smaller metabolic features that could give insights on metabolic heterogeneity within cell types. In the human fetal kidney, progenitor cells showed marked depletion of succinate in differentiated proximal tubules, consistent with earlier observations which relate stemness to higher succinate levels^17^. Fumarate, the downstream metabolite of succinate, shows a similar trend. We also observe a metabolic shift along nephron maturation characterized by a decrease in glutamine and a concomitant increase in glutamate (Supplementary Fig. 6).

Given the pronounced lipid specificity of proximal tubules, and our previous finding that fatty acid β-oxidation is critical for proximal tubule maturation^18^, we next examined lipid remodeling along the proximal tubule developmental trajectory. Spatial quantitative metabolomics using internal standards showed that developing proximal tubules accumulated higher levels of free fatty acids than other cell populations, with particularly strong enrichment of the long-chain polyunsaturated fatty acid docosahexaenoic acid (DHA) (Figure 2E). Analysis of signals related to phosphatidylethanolamine species further showed that proximal tubules were enriched in polyunsaturated phospholipids, including pre-annotated PE 38:4, PE 38:5, and PE 38:6, while containing relatively lower levels of more saturated species such as PE 38:1 (Figure 2F). This lipid composition is consistent with the specialized absorptive function of mature proximal tubules, which likely requires enhanced membrane fluidity^19^.

Notably, PE 38:6 was detected almost exclusively in mature proximal tubules (Figure 2G-H), and on-tissue MS/MS identified this species as containing DHA (Figure 2I). WNN-based pseudotime analysis further showed a progressive increase in PE 38:6 abundance along the trajectory from nephron progenitors to mature proximal tubules, indicating continuous metabolic remodeling during nephrogenesis (Figure 2J–K).

Together, these results show that polyunsaturated fatty acids, particularly DHA, progressively accumulate in developing proximal tubules and are incorporated into membrane phospholipids during maturation.

### DHA uptake programs accompany proximal tubule functionalization

DHA – and consequently PE 38:6 – accumulation may arise from increased cellular import, or endogenous biosynthesis. To investigate the source of DHA we turned to the transcriptional layer of our Stereo-meta-seq dataset. We first constructed a gene module consisting of *MFSD2A*, the canonical importer of LPC-DHA^20^, and *SLC27A1* to *SLC27A4* (*FATP1-4*), which function as FFA importers with acyl-CoA synthetase activity required for the incorporation of FFA into phospholipids. Visualization of the module scores in the developing kidney revealed a clear pattern. As cells mature, the DHA-import module score increased (Figure 3A–B). These findings were validated in an independent scRNA-seq dataset^21^ (Figure 3C), highlighting the robustness of transcriptional programs identified by Stereo-meta-seq.

**Figure 3.**
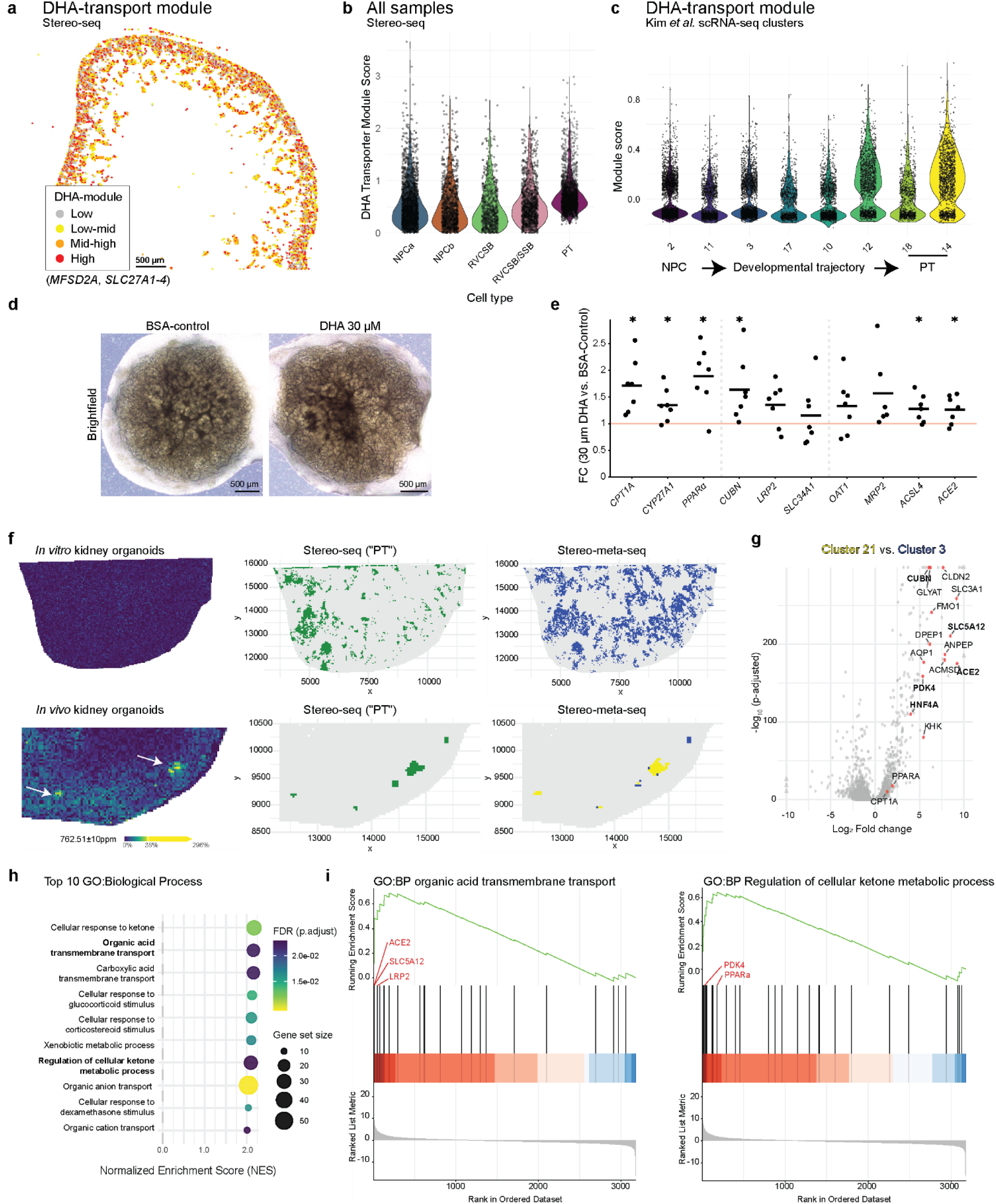
In vivo experiments show that PE 38:6 acquisition is associated with increased proximal tubule identity. (A) Module scores of DHA-transport genes in the human fetal kidney. (B) Quantification of module scores per cell type. (C) Independent validation of the same DHA-transport module using publicly available scRNA-seq^21^. (D) Representative brightfield images of BSA-control and DHA-treated kidney organoids. (E) qRT-PCR results of DHA-treated kidney organoids for the *PPARα* pathway, PT maturation and PT-markers. (F) In vivo transplantation of kidney organoids shows presence of PE 38:6 (left) in a population that is transcriptionally defined as “PT” in Stereo-seq (middle) and consists of two populations when clustered using WNN (right). (G) Volcano-plot showing an increase in expression of PT markers in the PE 38:6-associated cluster (yellow). (H) Gene Set Enrichment Analysis (GSEA) of the PE 38:6-positive proximal tubule cluster, highlighting the top 10 enriched biological processes. (I) Representative enrichment plots for two significantly enriched biological processes.

In our previous work, we found that kidney organoids exhibit a lipid metabolic state distinct from that of human fetal kidney^18^, which represents one barrier to their maturation. We therefore used Stereo-meta-seq to investigate the role of DHA and DHA-containing phospholipids, represented by PE 38:6, in iPSC-derived kidney organoids. In organoids maintained *in vitro*, of note DHA was not detected in the culture medium of kidney organoids.^18^ We next treated kidney organoids with DHA *in vitro* (Figure 3D) and found that DHA exposure increased the expression of proximal tubule markers and genes associated with epithelial maturation (Figure 3E). Previous studies have shown that activation of the PPARα pathway and enhanced fatty acid β-oxidation promote kidney organoid maturation^22^. Consistent with this model, DHA treatment modestly increased expression of PPARA and its downstream target CPT1A, suggesting that DHA promotes organoid maturation (Figure 3E).

We then examined kidney organoids after renal subcapsular transplantation, which in our previous studies produced a more mature developmental state than in vitro culture alone^23^. Importantly, transplanted organoids showed substantially greater DHA enrichment than organoids maintained in vitro (Figure 3F). PE 38:6 enrichment also emerged after transplantation, together with segregation of proximal tubules into PE 38:6-positive and PE 38:6-negative populations based on spatial multi-omics data (Figure 3F). PE 38:6-positive proximal tubules displayed activation of PPARα- and HNF4α-driven transcriptional programs and upregulation of functional maturation signatures, including organic acid transport pathways and ketone metabolism genes (Figure 3G–I). Together, these results suggest that DHA enhances proximal tubule maturation through multiple mechanisms, including activation of lipid-dependent transcriptional programs, such as PPARα signaling, and incorporation into membrane phospholipids during epithelial development. More broadly, the emergence of PE 38:6-positive and PE 38:6-negative proximal tubule states highlights the power of Stereo-meta-seq to resolve metabolite–transcript coupling underlying functional cell-state heterogeneity.

## Discussion

A central advance of this study is that it renders a longstanding question in developmental biology experimentally addressable: whether metabolic states merely accompany tissue maturation or actively instruct it. This conceptual shift is enabled by a technical one. By integrating spatial metabolomics and genome-wide transcriptomics within the same tissue section and at shared coordinates, Stereo-meta-seq allows metabolic and transcriptional states to be interrogated in direct spatial correspondence. Rather than adding a complementary layer of information, this approach exposes a previously inaccessible axis of tissue organization, making it possible to test whether local metabolic environments contribute causally to cell fate specification in situ.

Previous studies of kidney development have shown that metabolic remodeling accompanies the transition from nephron progenitors to differentiated epithelial cell types, with a pronounced shift toward fatty acid oxidation in the proximal tubule lineage ^18, 22^. These observations, together with differences between fetal tissue and organoids, have suggested that incomplete metabolic specification may constrain *in vitro* maturation. Here, we extend these findings beyond correlative trajectories by directly coupling metabolites and transcriptional programs in space. This reveals that the local lipid milieu, particularly DHA and its incorporation into membrane phospholipids, may contribute to its maturation.

Relative to existing same-section spatial multi-omics approaches^8–10^, Stereo-meta-seq provides several features that are critical for enabling these insights. On a single slide, Stereo-meta-seq combines high spatial resolution with genome-wide transcriptomic coverage, supports intrinsic spatial registration through MALDI ablation marks, and is compatible with multiple mass spectrometry platforms, enhancing robustness and scalability. Although limitations remain, including partial loss of transcriptomic signal following MSI and incomplete resolution of densely packed cellular regions, these do not obscure the central advance: the ability to resolve metabolite–gene relationships directly within intact tissue architecture.

This capability reveals an unanticipated aspect of nephrogenesis. Proximal tubule identity in the fetal kidney is not always robustly resolved by transcriptomic data alone, whereas lipid features markedly sharpen its definition as shown for proximal tubules.

This likely reflects an intrinsic property of developing tissues, in which cellular density, geometry, and mixing confound transcript-based annotation, while membrane composition more directly reports functional state. In this context, specific lipid species, including polyunsaturated phosphatidylethanolamines such as PE 38:6, provide a spatially stable readout of epithelial maturation that complements, and in some settings exceeds, the resolving power of transcript information.

Mechanistically, the data support a model in which DHA contributes to proximal tubule maturation through coordinated structural and regulatory effects. Incorporation of DHA into membrane phospholipids is expected to alter membrane fluidity^24^, curvature, and microdomain organization, consistent with the emergence of polarized epithelial architecture and brush border specialization^25, 26^. In parallel, in our data DHA is linked to activation of lipid-dependent transcriptional programs, including those governed by PPARα and HNF4α, which are central to fatty acid oxidation and proximal tubule identity^22^. Finally, progressive DHA enrichment may establish a metabolic state that supports the high energetic demand and limits stress susceptibility of mature proximal tubules. Consistent with this model, kidney organoids lack the DHA-associated lipid signature observed in fetal tissue, but acquire features of proximal tubule maturation following DHA supplementation or in vivo transplantation.

Together, these findings illustrate how acquisition of the Stereo-meta-seq modality can transform a descriptive framework into a mechanistic one by enabling direct spatial coupling of metabolites and gene expression and providing a framework for dissecting how local biochemical environments shape cell state in development and disease.

## Material & Methods

### Human samples

The collection and use of human material in this study were approved by the Medical Ethics Committee of the Leiden University Medical Center (P08.087). Gestational age was determined by ultrasonography in women undergoing elective pregnancy termination. Tissue from four embryos (15, 18, 19 and 21 weeks of gestation), was donated with written informed consent.

### Mouse samples

Kidney organoids transplantation experiments were approved by the Ethical Committee on Animal Care and Experimentation at Leiden University Medical Center (Permit No. AVD1160020186506). Animal experiments used for comparison of brass and plastic adaptors were approved by the Ethical Committee on Animal Care and Experimentation at Leiden University Medical Center (Permit No. AVD1160020172926). All work with animals was performed in compliance with Dutch government guidelines.

Mouse kidney experiments for multi-omics measurement (n=3) were approved by the Animal Ethics Committee of Fudan University, and performed in accordance with institutional guidelines (protocol number 2023-IBS-01JZS). Mouse brain experiments (n=3) were approved by the Institutional Animal Care and Use Committee of Westlake University and performed in accordance with the protocol AP#24-112-ZYL. Mouse liver experiments conducted for MALDI-TOF were performed in accordance with the guidelines of the Institutional Animal Care and Use Committee at the Shanghai Institute of Biochemistry and Cell Biology (protocol number SIBCB-NAF-14-001-S327-010).

### Conductive adapter design and fabrication

The conductive adapter was computer numerical control (CNC)-machined from brass to overall dimensions of 75 × 25 × 1.1 mm (length × width × thickness). Four central recesses (1 × 1 cm, approximately 0.71 mm depth) were milled to securely hold Stereo-seq chips, featuring corner extraction holes to facilitate chip removal post-MSI. A 6.61 mm margin was maintained between the peripheral recesses and the adapter edge.

Crucially, the recess depth was strictly matched to the chip height (tolerance < 10 µm). This precision engineering ensured that the surfaces of the brass adapter and the mounted chip were perfectly aligned on the same focal plane, which is essential for accurate metabolomic detection. The non-conductive plastic adaptor was fabricated with PMMA.

### Validation of conductive adapter

Mouse liver experiments conducted for MALDI-TOF were performed in accordance with the guidelines of the Institutional Animal Care and Use Committee at the Shanghai Institute of Biochemistry and Cell Biology. The approved animal protocol numbers are SIBCB-NAF-14-001-S327-010. Mouse brain experiments were approved by the Institutional Animal Care and Use Committee of Westlake University and performed in accordance with the protocol AP#24-112-ZYL.

### RapifleX MALDI-MSI

Liver tissues embedded in 25% gelatin were cryosectioned (16 µm thickness) at −20 °C using a Cryostar NX70 cryostat (Thermo Fisher Scientific). Sections were thaw-mounted onto ITO-coated glass slides or Stereo-seq chips, with the latter subsequently seated in the conductive brass adapter. Mounted sections were vacuum freeze-dried for 15 min. A MALDI matrix solution of 7 mg/mL N-(1-naphthyl) ethylenediamine dihydrochloride (NEDC; Sigma-Aldrich) in ethanol/acetonitrile/water (70:25:5, v/v/v) was applied using an HTX TM-Sprayer (HTX Technologies). Spraying parameters were set as follows: flow rate 0.05 mL/min, nozzle velocity 1,200 mm/min, nozzle temperature 70 °C, nozzle height 40 mm, and drying gas pressure 10 psi, with 30 passes and a 30 s drying time between passes.

MALDI-TOF MSI was performed using a RapifleX system (Bruker Daltonics) calibrated with red phosphorus. Negative ion-mode mass spectra were acquired across a mass range of *m/z* 80–1000 at a 20 × 20 µm² pixel size, using 200 laser shots per pixel at a 5 kHz laser repetition rate. Data acquisition and visualization were performed using flexControl 4.2 and flexImaging 6.0 (Bruker Daltonics). Following the initial MSI acquisition on the brass adapter, the Stereo-seq chips were transferred to a plastic adapter for repeat MSI detection. Downstream data analysis was conducted using SCiLS Lab (Bruker) and mMass.

### Quantitative Mass Spectrometry Imaging (qMSI)

16 µm thick tissue sections were cryosectioned using a Cryostar NX70 cryostat (Thermo Fisher Scientific) at −20 °C and placed directly onto Stereo-seq T-chips v1.3 (STOmics, MGI, Latvia) using the cold-mounting method as described by the manufacturer. To preserve metabolites, chip warming was reduced to a minimum by only placing a finger on the back of the chip.

For qMSI, U-¹³C-labelled metabolite yeast internal standard (99%; ISO1, Cambridge Isotope Laboratories), followed by NEDC matrix were applied as previously described^14^. Data were acquired using a TimsTOF fleX MALDI-2 system (Bruker Daltonics) in negative mode with MALDI2 at the mass range of *m*/*z* 50–1,000. The MALDI-2 laser had a pulse delay time of 10 μs. Data were acquired at 16 x 16 µm pixel size with a beam scan area of 20 x 20 µm. The laser was operated at 1 kHz with 100 laser shots accumulated per pixel. Data acquisition was performed using flexControl (v.4.0, Bruker Daltonics).

### Tandem Mass Spectrometric (MS/MS) analysis

Image-guided MS/MS was performed on a tissue section of human fetal kidney sprayed with NEDC matrix. Regions of interest containing proximal tubules, as delineated by presence of *m/z* 762.51 were determined using a consecutive slide. MS/MS was performed on *m/z* 762.51 using a MALDI-2-timsTOF fleX instrument in MALDI-2 mode. Negative ion mode spectra with a mass range of *m/z* 50-1000 were acquired with beam scan mode on 46 µm by 46 µm, with laser power at 50% and 250 shots per pixel. To fragment *m/z* 762.51, an isolation window of *m/z* 8 was selected with a 40 eV collision energy.

### Stereo-seq

Immediately after qMSI data acquisition, the NEDC matrix on the chips was removed with cold methanol for 10 seconds. Upon continuation of the protocol, chips were processed according to the manufacturer’s guidelines, beginning with the −20°C methanol fixation step.

### Kidney organoid culture and DHA treatment

For transplantation experiments, human induced pluripotent stem cells (hiPSC-MAFB^27^) were maintained on vitronectin-coated plates in Essential 8 medium (Thermo Fisher Scientific) and passaged using 0.5 mM Ultra-Pure EDTA (Thermo Fisher Scientific) at 37°C in a humidified atmosphere containing 5% CO₂. One day prior to initiation of differentiation, cells were dissociated using TrypLE Select (Thermo Fisher Scientific) and replated as single cells in E8 medium (Thermo Fisher Scientific) supplemented with RevitaCell (Thermo Fisher Scientific). Differentiation to kidney organoids was performed following previously established protocols^23, 28^. Cells were cultured for 4 days in STEMdiff APEL-2 medium (Stem Cell Technologies) supplemented with 1% PFHMII (Life Technologies) and 1% Antibiotic-Antimycotic (Life Technologies) and differentiation was initiated with 8 μM CHIR99021 (Tocris). On day 4, the medium was replaced with APEL-2 containing 200 ng mL⁻¹ recombinant human FGF9 (R&D Systems) and 1 μg mL⁻¹ heparin (Sigma-Aldrich). On day 7, cells were exposed to a 1-hour pulse of 5 μM CHIR99021, dissociated, and transferred to 3D culture on polyester transwell membranes (0.4 μm pore size, Corning). After 5 days, growth factors were withdrawn, and medium changes with APEL-2 were performed every 2 days. Organoids were collected and snap frozen at day 7+18.

For experiments with DHA, hiPSC-MAFB was cultured in mTESR Plus (Stem Cell Technologies) on human recombinant laminin-521 (BioLaminin) and passaged using 0.5 mM EDTA. For differentiation, cells were plated as single cells on laminin in mTESR Plus containing 10 µM Y-27632 dihydrochloride (ROCK inhibitor, Tocris) and the next day differentiation was initiated using 6 µM CHIR99021 in TeSR-E6 medium (E6, Stem Cell Technologies) supplemented 1% Antibiotic-Antimycotic (Life Technologies). From day 4 until day 7+14, the differentiation proceeded as described above in E6 medium. From day 7+14 to day 7+18, organoids were treated with either 30 µM BSA-Docosahexaenoate Polyunsaturated Fatty Acid Complex (Cayman Chemical) and 30 µM BSA Control for BSA-Fatty Acid Complexes (Cayman Chemical) in E6. Seven independent batches of kidney organoids were generated and analyzed Using quantitative real-time PCR.

### Kidney organoid transplantation

All animal experimental protocols were approved by the animal welfare committee of the Leiden University Medical Center. Recipient mice (n=5, 8 weeks old NOD/SCID, Charles River Laboratories) received buprenorphine as pre- and postoperative analgesic in water and subcutaneously and were anesthetized with isoflurane. Core body temperature was maintained at 37 °C. Via flank incisions, kidneys were exteriorized and kidney organoids, cultured for 7+18 days were bisected and transplanted under renal capsule in the left and right kidney. The mice were sacrificed after 28 days and the collected kidneys were snap frozen.

### RNA isolation and quantitative real-time PCR

Total RNA was extracted from 2 kidney organoids from each experiment using the Maxwell RSC simplyRNA Cells Kit (Promega) on the Maxwell RSC 48 Instrument (Promega) according to the manufacturer’s protocol. RNA concentration and purity were assessed by NanoDrop spectrophotometer (Thermo Fisher Scientific). cDNA was synthesized from 500 ng total RNA using iScript Reverse Transcription Supermix (Bio-Rad) in a 20 μL reaction volume (priming at 25°C for 5 min, reverse transcription at 46°C for 20 min, and RT inactivation at 95°C for 1 min). cDNA was diluted 1:5 in nuclease-free water before use as qPCR template.

Quantitative real-time PCR was performed using iQ SYBR Green Supermix (Bio-Rad) on the CFX Connect Real-Time System (Bio-Rad). Each reaction contained 1 μL diluted cDNA template, 500 nM forward and reverse primers, and 1× iQ SYBR Green Supermix in a total volume of 10 μL (protocol: denaturation at 95°C for 3 min, followed by 40 cycles of 95°C for 10 sec and 65°C for 30 sec). All reactions were run in triplicate. Primer sequences are listed in **Supplementary Table 1**.

Gene expression levels were normalized to 18S ribosomal RNA (RN18S) as the reference gene. For each independent differentiation experiment, fold change was calculated using the ΔΔCt method relative to the matched BSA vehicle control within the same batch. Fold changes from independent experiments (n = 7 biological replicates from independent differentiations) were log₂-transformed and analyzed by one-sample t-test. P < 0.05 was considered statistically significant.

### Modality-matching

Using cellbin2’s matrix.py (https://github.com/STOmics/cellbin2/tree/dev), a .tiff of the Stereo-seq transcriptomics was generated. At least three laser-track landmarks were identified (Figure 1, box 3). Subsequently, qMSI coordinates were transformed using an affine-transformation. Next, qMSI pixels were matched based on a nearest neighbor match of the transformed coordinates onto the Stereo-seq object binned to the same size as the qMSI beam scan area on a one-to-one basis and distances to the nearest neighbor visualized spatially and checked for accurate mapping.

### Stereo-seq data analysis

Raw sequencing data was aligned using Stereo-seq Analysis Workflow (SAW) 8.1. Data was loaded into RStudio Server (version 1.3.959, R version 4.4.1) and placed in a Seurat v5 object (version 5.3.0)^29^. Mitochondrial and ribosomal genes were filtered out.

Samples were harmonized using Stereopy (version 1.6.0)^30^, which enabled calculation of the PCA based on the complete transcriptome. Empty bins were filtered out. Data was normalized using total-count scaling followed log-transformation. Neighborhood graphs were constructed using the integrated Principal Components (PCs) (50 dim, k = 8) followed by UMAP for dimensionality reduction and clustering using the Leiden algorithm at resolution 1.0. All downstream analysis was performed in RStudio Server.

### qMSI data analysis

Peak picking for clustering (*m/z* > 400) was performed on the average spectrum by initial selection based on relative intensity (>0.01%) in mMass after which manual curation of selected peaks took place. Matrix features were excluded from further analysis. Total Ion Count intensities were exported for clustering.

For metabolite feature annotation, a previously generated^14^ list of *m/z* features was used for which the U-^12^C and U-^13^C variants of the metabolite were identified within the dataset. Metabolites were normalized per pixel to their U-^13^C counterpart.

### Integrated analysis (WNN)

The first 20 PCs of both Stereo-seq and qMSI RMS normalized features were used as input for Seurat’s FindMultiModalNeighbors, after which UMAP was performed and WNN clusters were calculated.

### Cell type annotation

Identified cell clusters were manually annotated based on their marker gene expression as identified in single cell RNA-seq (scRNA)^31^, together with their spatial localization within the tissue, location by Uniform Manifold Approximation and Projection (UMAP) and lipid markers. Populations that showed a mixture of two or multiple cell types due to the resolution have been classified as such, for example RVCSB (Renal Vesicle, Comma Shaped Body). Some WNN-clusters were subclustered to reveal additional populations.

### Pseudotime analysis

To construct a pseudotime trajectory of proximal tubule development cell types known to give rise to proximal tubules were selected. From these cell types, diffusion components (DCs) were calculated using destiny (version 3.18.0). DC1 represented the developmental trajectory in pseudotime.

### Differential expression

Differences in transcriptomic and lipidomic profiles between identified cell clusters were identified using Seurat’s *FindMarkers* function.

### Data visualization

Data was visualized using ggplot2 (version 4.0.1).

### Data availability

Raw and processed Stereo-seq sequencing data is available from EGA and ENA under accession numbers **XXX.** Additional data files and Seurat objects are available at Zenodo **XXX**. Raw qMSI data related to this study will be made available upon reasonable request to the corresponding authors due to the size these datasets. The code used in this study is available at GitHub (**LINK**)

---

**Supplementary Fig. 1.**
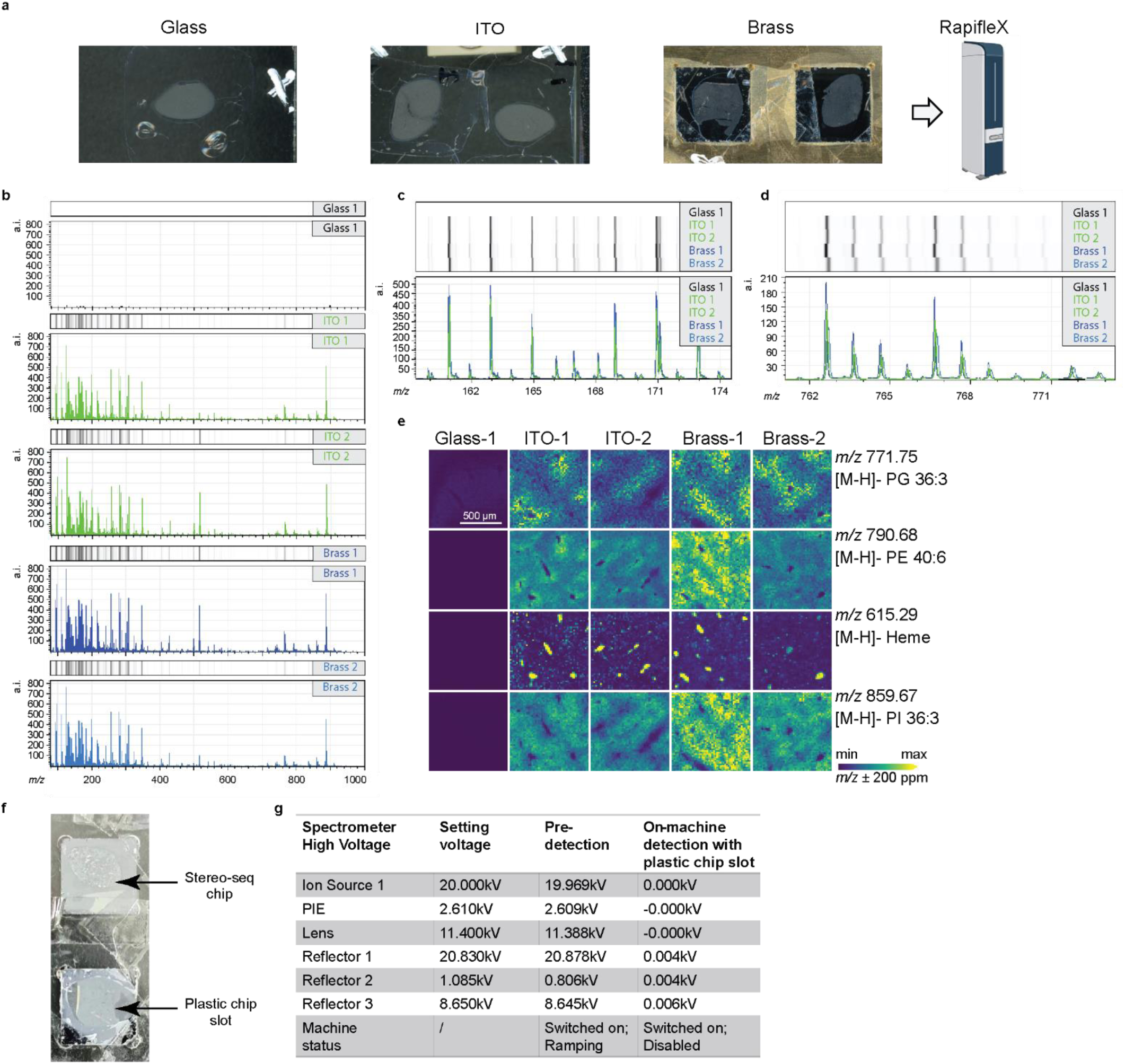
Validation of MALDI-TOF MSI performance using conductive adapters. (A) Experimental design comparing MALDI-TOF signal acquisition across non-conductive glass slides (Glass), ITO-coated slides (ITO), and Stereo-seq chips mounted on brass adapters (Brass) using mouse liver sections. (B) Comparison of average mass spectrometry signal intensities across the three materials (Glass, n=1; ITO and Brass, n=2), m/z 80–1000. (C-D) Magnified views of the average mass spectra for selected low-mass (*m/z* 160–175 in C) and high-mass (*m/z* 760–775 in D) ranges. (E) Spatial distribution of representative liver metabolites across the three slide types. (F) Schematic illustration of MALDI-TOF detection utilizing a non-conductive plastic adapter for Stereo-seq chips. (G) Voltage monitoring during acquisition with the plastic adapter, showing the set voltage, pre-detection standby state, and actual operational voltage during scanning.

**Supplementary Fig. 2.**
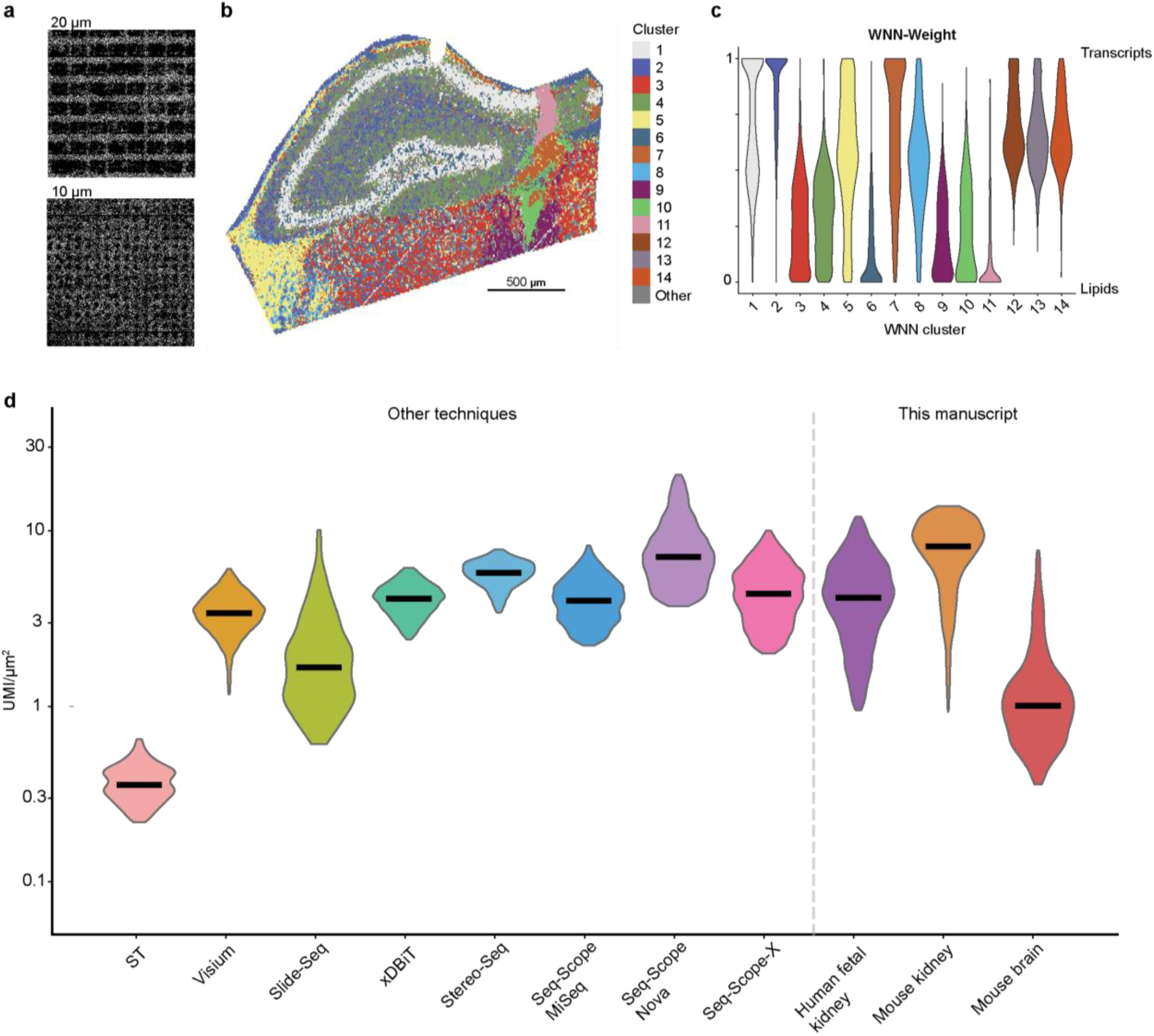
Stereo-meta-seq can be applied to different tissues with a resolution up to 10 µm. (A) Representative image of the fiducial laser ablation marks at 20 µm and 10 µm. (B) WNN based clustering of the mouse hippocampus recapitulating the major anatomical regions. (C) Associated distribution of WNN-weights per cluster. (D) Comparison of UMI/µm^2^ of single-omics techniques in comparison to Stereo-meta-seq. Part of the figure was adapted from a previous report^15^. All datasets subsampled to n=5,000 with P1–P99 clipping for visual consistency.

**Supplementary Fig. 3.**
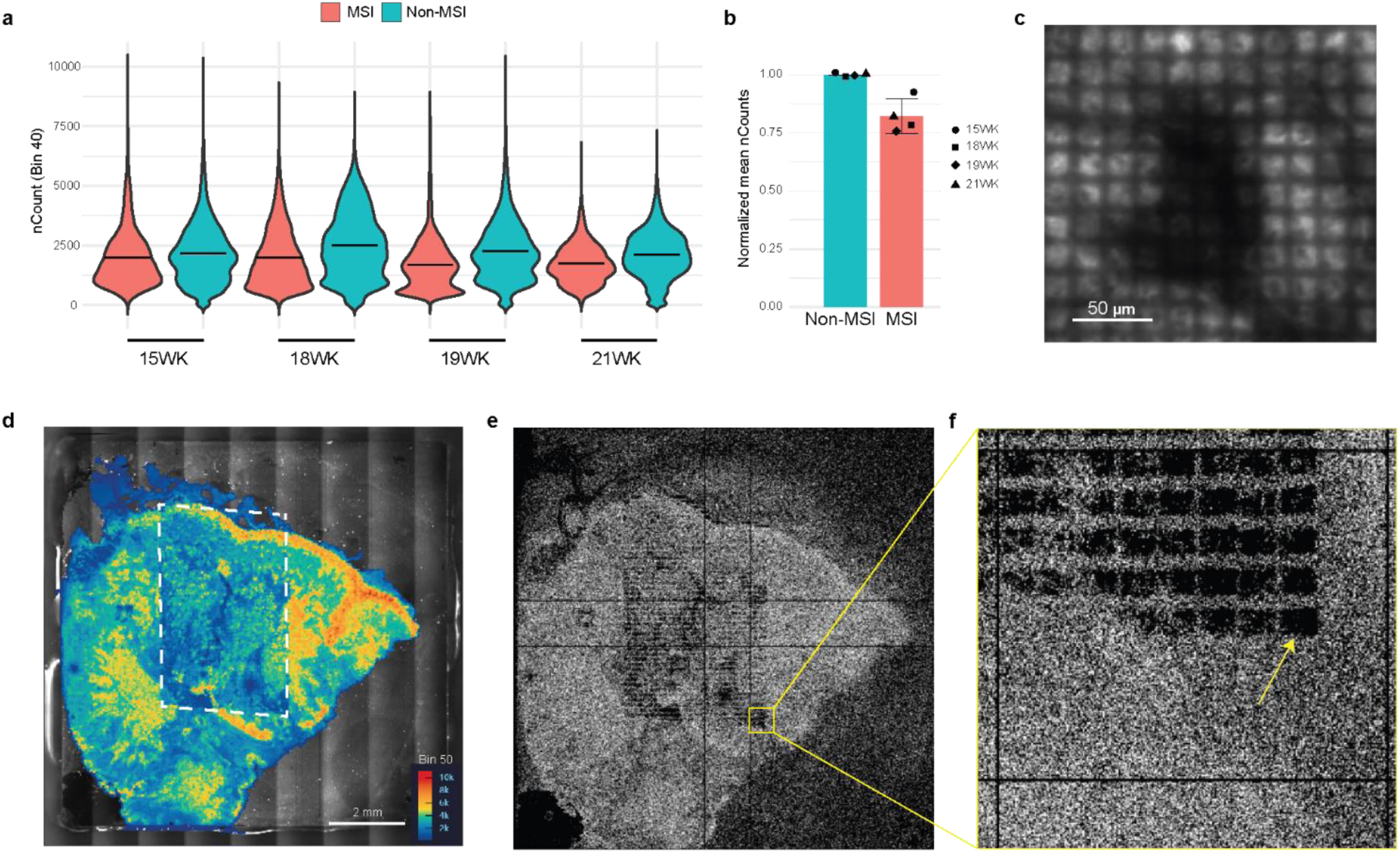
Laser ablation by qMSI leaves fiducial marks but minimally affects transcriptomic Stereo-seq readout. (A) Quantification of nCount per bin40 for all samples in non-MSI (Stereo-seq-only) and MSI areas. (B) Quantification of all replicates. (C) Autofluorescence imaging showing laser marks in the tissue after qMSI (D) STOmics’ StereoMAP visualization of nCount of one representative sample. White box indicates area of qMSI measurements. (E) Cellbin2’s matrix.py output depicting 1 bin per pixel, showing laser ablation marks. (F) Zoom-in, yellow arrow indicates a single 16 µm measurement. Vertical and horizontal lines are Stereo-seq track-lines.

**Supplementary Fig. 4.**
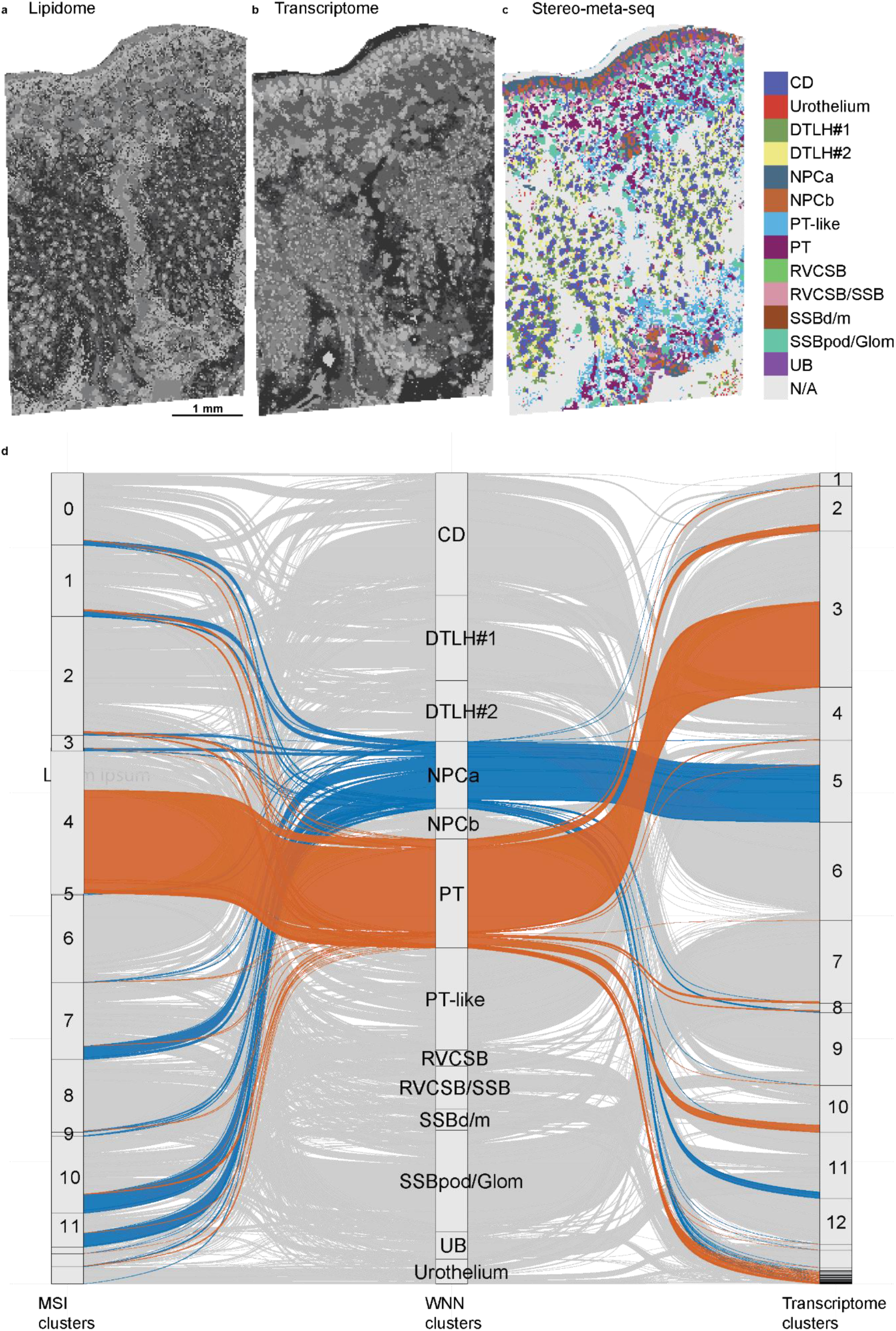
Combining the modalities of Stereo-meta-seq leads to more clearly defined clusters than individual modalities. (A) Cell populations as defined by Stereo-seq data alone. The nephron progenitor (NPC) zone is clearly defined. (B) Cell populations as defined by the qMSI lipidome alone. The collecting duct (CD) and proximal tubule (PT) areas are clearly defined. (C) Definitive WNN-based clustering. NPC, CD and PT areas are well defined. (D) Alluvial plot showing the final WNN-clusters and the distribution of NPCa (blue) and PT (orange) cell types in individual modalities.

**Supplementary Fig. 5.**
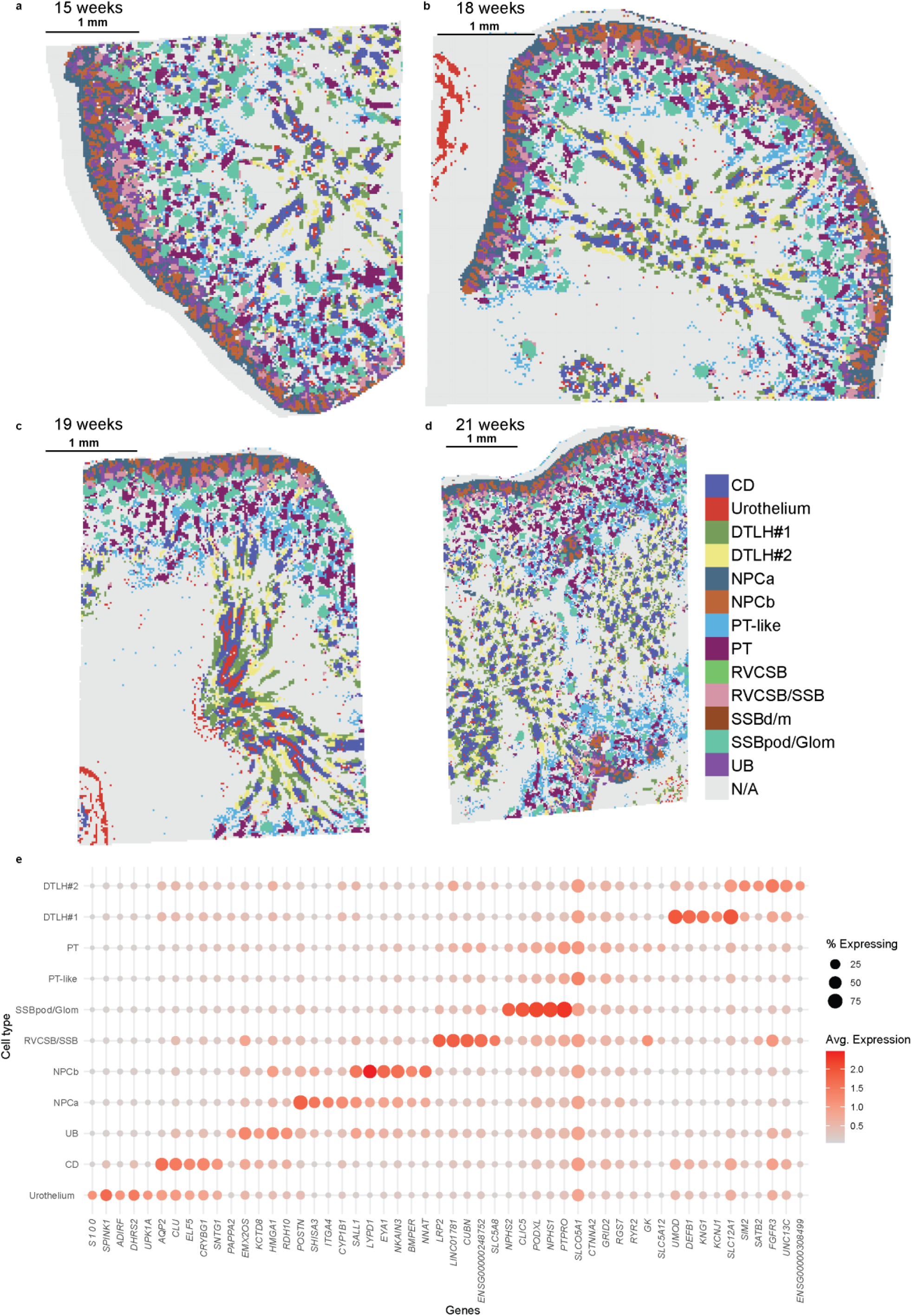
Stereo-meta-seq enables detailed cell type annotation recapitulating fetal kidney anatomy. (A-D) Spatial visualization of all samples of human fetal kidneys, ranging from gestational ages of 15 to 21 weeks. (E) Dotplot showing 5 marker genes per identified cell populations as defined by Seurat’s *FindAllMarkers*.

**Supplementary Fig. 6.**
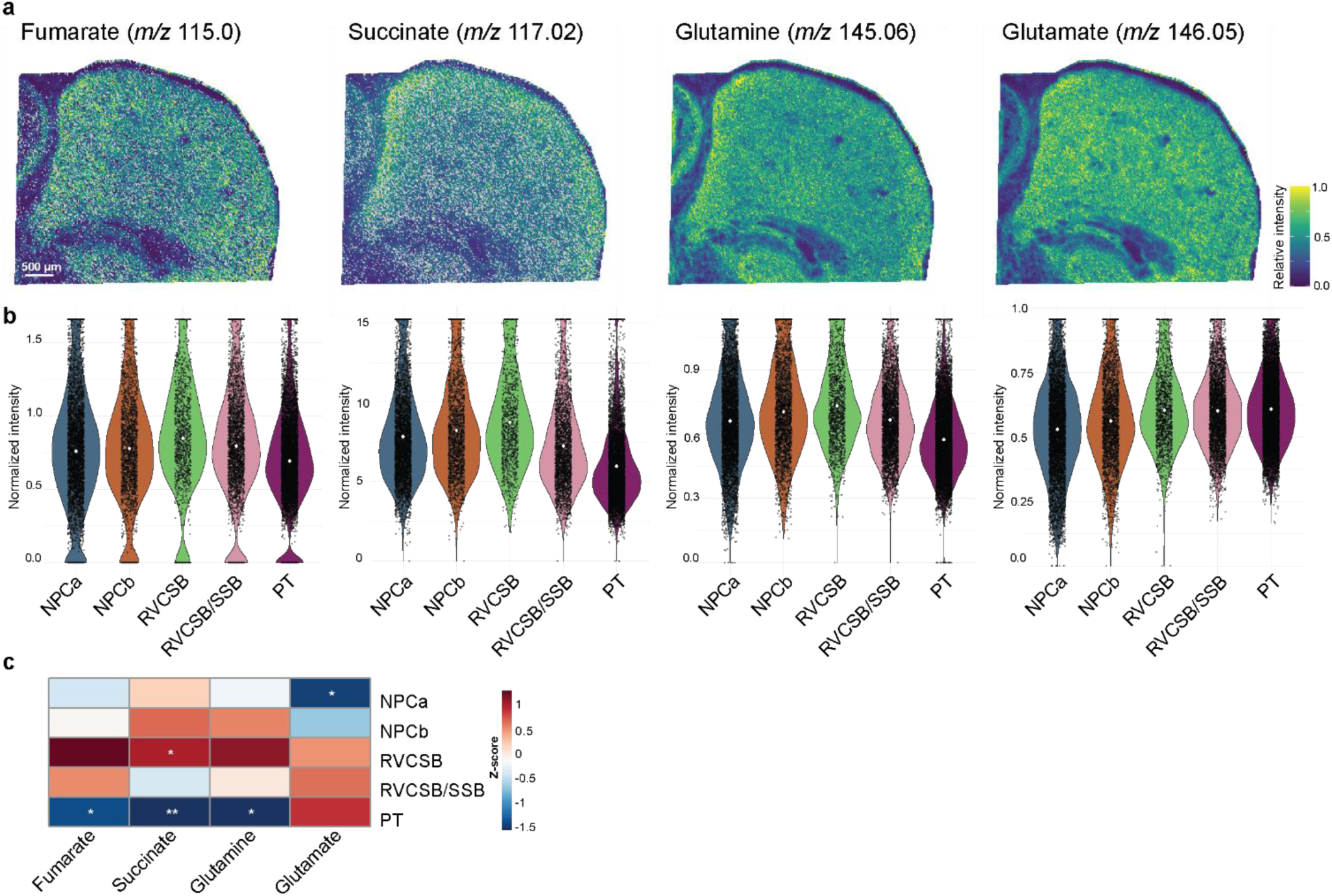
Stereo-meta-seq provides normalized data for small metabolic features. (A) Spatial visualization of the metabolites fumarate, succinate, glutamine and glutamate in the human fetal kidney. (B) Quantification of the normalized intensities for the developmental trajectory from NPC towards PT. (C) Heatmap showing Z-scored mean metabolite intensities across annotated cell types. Pixel intensities were pseudobulked per biological sample prior to statistical testing. Significance indicates enrichment or depletion relative to other cell types across samples (* = *p* < 0.05, ** = *p* < 0.01; *n* = 4 samples; BH-corrected)

**Table S1.**
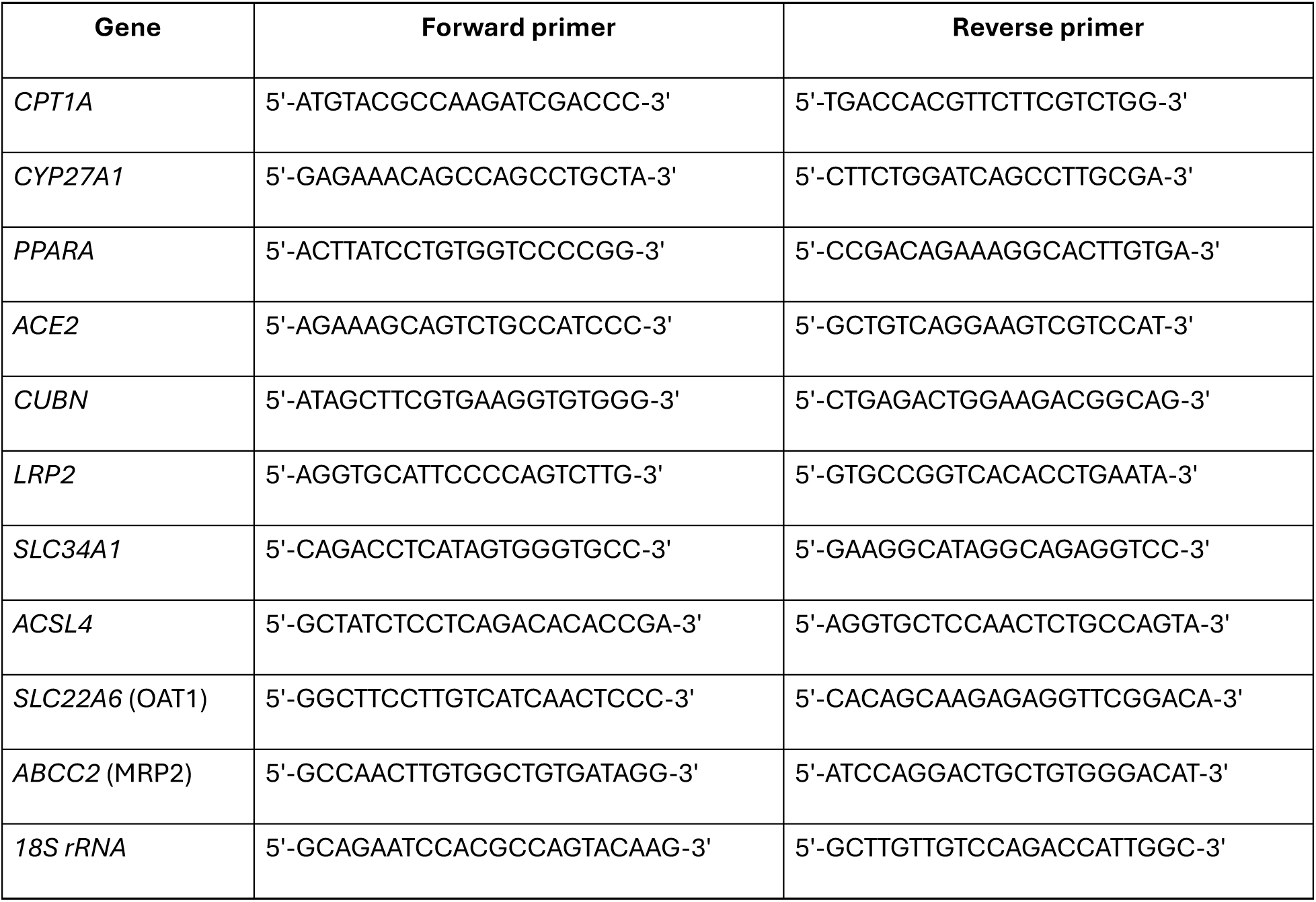
Primer sequences used for quantitative real-time PCR.

## Contributions

R.H.v.S. processed the human fetal kidney, brain and kidney organoids data, developed the scripts, made figures and wrote the manuscript. B.W. invented the brass adapter and performed the rapifleX experiments. Y.Zh. performed the mouse kidney experiments, organoid experiments and processed the mouse kidney data. S.S. processed the human fetal kidney data, annotated clusters and developed the associated scripts. F.W. and Y.Zou performed the mouse brain experiments. D.S provided valuable input on DHA-metabolism. I.S. performed the On-tissue MS/MS measurements. C.W.v.d.B supervised organoid work and conducted the transplantation experiments. B.M.v.d.B., U.W. and M.G. provided feedback on the manuscript. G.W. conceptualized and developed Stereo-meta-seq, conducted experiments, and provided feedback on the manuscript and secured funding. T.J.R. secured funding, provided supervision, and reviewed the manuscript. L.H. secured funding, provided supervision, and reviewed the manuscript.

## Acknowledgements

We gratefully acknowledge Manon Zuurmond (Department of Internal Medicine, LUMC, Leiden, The Netherlands) for making the illustration shown in Figure 1A. We thank Ellen Lievers, Rianne Y. van Nieuwland, Anneloes H. Verwey for technical assistance. We also express our gratitude to members of the Rabelink Lab for their constructive feedback on the manuscript. We acknowledge Ao Chen, Miguel A Esteban, Yiwei Lai from BGI research and Jia Khoo, Li Pu from BGI Europe for their support on Stereo-seq experiments. We thank the Shanghai Institute of Nutrition and Health for providing support with the MALDI-TOF measurements. The Novo Nordisk Foundation Center for Stem Cell Medicine (reNEW) is supported by Novo Nordisk Foundation grant (NNF21CC0073729). Ton J. Rabelink is funded by the European Union through ERC grant (SPARK 101140863). This study was supported by the National Natural Science Foundation of China (92368301, U25A600024) to Lijian Hui and Gangqi Wang. Lijian Hui was supported by the Noncommunicable chronic disease-national science and technology major project (2024ZD525404), the National Key Research and Development Project (2024YFA1803101, 2024YFA1803303), the Shanghai Municipal Science and Technology Major Project. Yilong Zou was supported by the National Natural Science Foundation of China (32450793).

